# High resolution mapping of novel non-transgressive hybrid susceptibility in barley exploited by *P. teres* f. *maculata* maps to a single pentatricopeptide repeat-containing protein

**DOI:** 10.1101/2024.03.17.585425

**Authors:** Shaun J. Clare, Abdullah F. Alhashel, Mengyuan Li, Karl M. Effertz, Roshan Sharma Poudel, Jianwei Zhang, Robert S. Brueggeman

**Affiliations:** Department of Crop and Soil Sciences, Washington State University, Pullman, WA 99164, USA; Department of Plant Pathology, North Dakota State University, Fargo, ND 58108-6050, USA; Department of Plant Protection, College of Food and Agriculture Sciences, King Saud University, Riyadh 11451, Saudi Arabia; National Key Laboratory of Crop Genetic Improvement, Huazhong Agricultural University, Wuhan 430070, China; Dewey Scientific, Pullman, WA 99163, USA; Syngenta Seed Inc., Durham, NC 27709, USA

## Abstract

Hybrid genotypes can provide significant yield gains over conventional inbred varieties due to heterosis or hybrid vigor. However, hybrids can also display unintended negative attributes or phenotypes such as extreme pathogen susceptibility. The necrotrophic pathogen *Pyrenophora teres* f. *maculata* (*Ptm*) causes spot form net blotch, which has caused significant losses to barley worldwide. Here, we report on a non-transgressive hybrid susceptibility locus in barley initially recognized because the three parental lines CI5791, Tifang and Golden Promise are resistant to *Ptm* isolate 13IM.3, however F_2_ progeny from CI5791 × Tifang and CI5791 × Golden Promise crosses exhibited extreme susceptibility. The susceptible phenotype segregated in a ratio of 1 resistant:1 susceptible representing a genetic segregation ratio of 1 parental (res):2 heterozygous (sus):1 parental (res) suggesting a single hybrid susceptibility locus. Genetic mapping using a total of 715 CI5791 × Tifang F_2_ individuals (1430 recombinant gametes) and 149 targeted SNPs delimited the hybrid susceptibility locus designated *Susceptibility to Pyrenophora teres 2* (*Spt2*) to an ∼198 kb region on chromosome 5H of the Morex V3 reference assembly. This single locus was independently mapped with 83 CI5791 × Golden Promise F_2_ individuals (166 recombinant gametes) and 180 genome wide SNPs that colocalized to the same *Spt2* locus. The CI5791 genome was sequenced using PacBio Continuous Long Read technology and comparative analysis between CI5791 and the publicly available Golden Promise genome assembly determined that the delimited region contained a single high confidence *Spt2* candidate gene predicted to encode a pentatricopeptide repeat-containing protein.

## Introduction

Hybrid barley was one of the first commercially available small grain hybrid crops in the world and has been grown since the 1980s [1]. Currently, hybrid winter feed barley is grown on more than 200,000 ha in Europe [1]. Thus, hybrid barley may also be a solution to boost malt barley yields to satisfy the demands of the brewing and distilling industries as climate change is predicted to cause world malt barley shortages [2, 3]. Models show that some barley growing regions will experience extreme heat and drought while other production regions will experience increased temperatures and excess moisture contributing to conditions conducive to disease development impacting yield and quality [3]. For some host-pathogen genetic interactions, the use of hybrid barley may increase disease resistance [4], however in other pathosystems, hybrids may reduce resistance or increase susceptibility. The outcome of these interactions also depends on the virulence mechanisms of the pathogen and the nature of the pathogen in terms of acquiring nutrients from the host, either as biotrophic or necrotrophic pathogens.

Plant-pathogen genetic interactions that determine compatible (susceptible) -vs- incompatible (resistant) interactions, typically rely on resistance (*R*) genes that encode receptor-like proteins/kinases (RLPs/RLKs) or nucleotide binding-leucine rich repeat (NLR) class proteins that recognize pathogen effectors or their action [5]. However, diverse coevolution that occurs in pathosystems can also evolve into non-dogmatic interactions that do not conform to this typical mechanism such as the first cloned resistance gene, *Hm1*, which encodes a reductase that neutralizes a toxin [6]. Additionally, increased characterization of resistance genes has identified novel *R*-protein classes that may or may not include integrated domains that follow the proposed integrated sensory or decoy domain model such as ankyrin-repeat domains or WRKY domains [7–10]. WRKY transcription factors have already been implicated in resistance with *HvWRKY6* required for *Pyrenophora teres* f. *teres* (*Ptt*) resistance in barley [11]. Therefore, these studies show that alternative mechanisms and classes of genes/proteins are involved in plant resistance responses and non-canonical candidate genes should be thoroughly investigated.

In addition, susceptibility genes and their respective proteins are manipulated by necrotrophic fungal pathogens to facilitate disease development and have often initially been characterized as recessive resistance genes. The first confirmed necrotrophic pathogen susceptibility gene identified was *Pc-2/Vb* conferring dominant resistance to oat crown rust and dominant susceptibility to Victoria blight (Lorang *et al.* 2007). This was the first study showing that necrotrophic pathogens can hijack dominant resistance genes to elicit programmed cell death (PCD) responses, referred to as the hypersensitive response (HR). These PCD/HR responses provide effective resistance against biotrophic pathogens that require living host tissue to acquire nutrients by sequestering the biotrophic pathogen to small foci of dead cells. However, when these mechanisms are elicited by necrotrophic effectors (NEs) secreted by necrotrophic pathogens that require dead or dying tissue to acquire nutrients, the PCD response results in host susceptibility and continued pathogen proliferation [12].

In the barley-*Ptt* necrotrophic pathosystem (causal agent of net from net blotch), the dominant susceptibility locus *Spt 1*was first identified as the recessive resistance loci *rpt.r* and *rpt.k* in Rike and Kombar, respectively [13, 14]. The *Spt1* locus was hypothesized to be two tightly linked genes providing recessive resistance to different *Ptt* isolates. However, high-resolution mapping led to the hypothesis of a single gene allelic series that confers isolate specific *Spt1*-mediated susceptibility [15, 16]. In addition, alleles of this same gene may also correspond to the dominant *Rpt5-*mediated broad-spectrum resistance that mapped to the same region in CI5791 [16, 17]. These types of complex genetic interactions of susceptibility and/or resistance genes could have relevant effects on disease development when hybrids are deployed depending on the combinations of alleles and effector repertoires of pathogen isolates challenging the plant.

*Pyrenophora teres* f. *maculata* (*Ptm*) recently diverged from the close relative *Ptt* and is the causal agent of the barley disease spot form net blotch (SFNB) [18, 19]. SFNB is present worldwide and can cause significant yield losses, typically between 10-40%, but can also result in complete crop failure or degraded seed quality [20–22]. To date, over 150 quantitative trait loci (QTL) have been mapped within the host of the barley-*Ptm* pathosystem [16] and seven regions within the wheat-*Ptm* pathosystem [23]. Here, we report on the high-resolution mapping of a non-transgressive hybrid susceptibility gene designated *Susceptibility to Pyrenophora teres 2* (*Spt2*) in response to infection from *Ptm* isolate 13IM8.3. The *Ptm* isolate 13IM8.3, is avirulent on CI5791 and Tifang, is hyper virulent on all CI5791 × Tifang F_1_ individuals, and was used to genetically characterize the *Spt2* hybrid susceptibility locus (**Figure 1**). Thus, a CI5791 × Tifang F_2_ population was utilized for high-resolution mapping of the *Spt2* locus to an ∼198 kb region containing a single high confidence gene model predicted to encode a pentatricopeptide repeat-containing protein based on the cultivar Morex V3 genome assembly. To our knowledge, the *Spt2* locus is the first reported case of hybrid susceptibility within the same species, as opposed to hybridization of two closely related species. The *Spt2-*mediated susceptibility to *Ptm* isolate 13IM8.3 is non-transgressive, similar to heterosis, as only heterozygous individuals at the *Spt2* locus are susceptible and is therefore not genetically stable [24]. This phenomenon is also not the result of post-zygotic genetic incompatibilities such as hybrid necrosis that involves epistatic interaction between multiple loci and occurs in the absence of the pathogen [25, 26].

**Figure 1.**
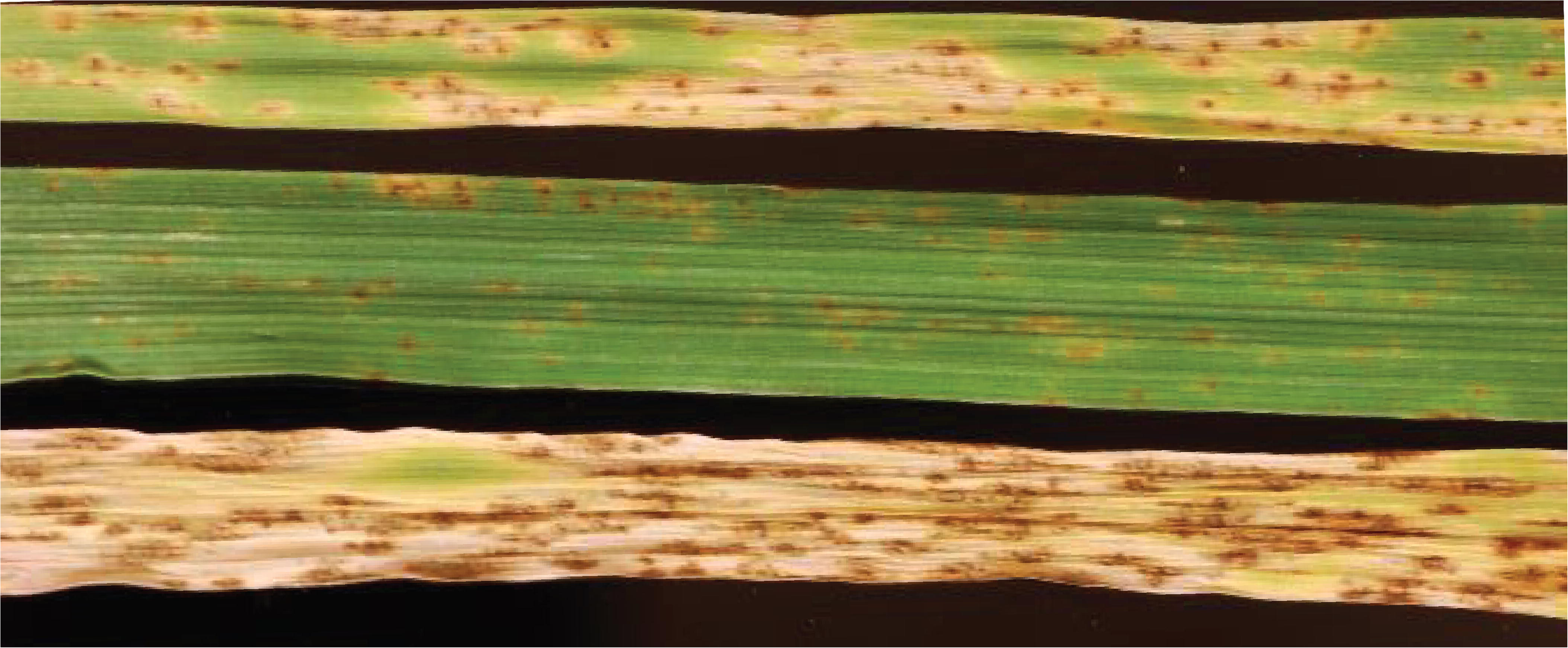
Phenotypic results of parental and hybrid barley. Parents CI5791 (top), and Tifang (middle) and a CI5791 × Tifang F1 after one week post inoculation with *Pyrenophora teres* f. *maculata* isolate 13IM8.3.

## Results

### Initial CI5791 × Tifang Genetic Mapping with SNP Markers Derived from the 9K SNP Array

The initial genetic mapping of *Spt2* was conducted with the phenotyping of 106 CI5791 × Tifang F_2_ individuals that were genotyped with a 365 PCR-GBS SNP marker panel resulting in 119 high quality polymorphic SNP markers spread across the barley genome. The QTL mapping detected a single highly significant locus on chromosome 5H delimited by the markers 11_10641 and 11_20236 with the most significant marker 12_20350 giving a LOD score of 23.1 (**Figure 2A**). LOD thresholds were calculated at 3.788 (α = 0.05) and 4.701 (α = 0.01). The 16.7 cM genetic interval (**Figure 3A**) corresponded to 23.1 Mb of genome sequence from the Morex V3 reference genome assembly containing 285 annotated genes [27].

**Figure 2.**
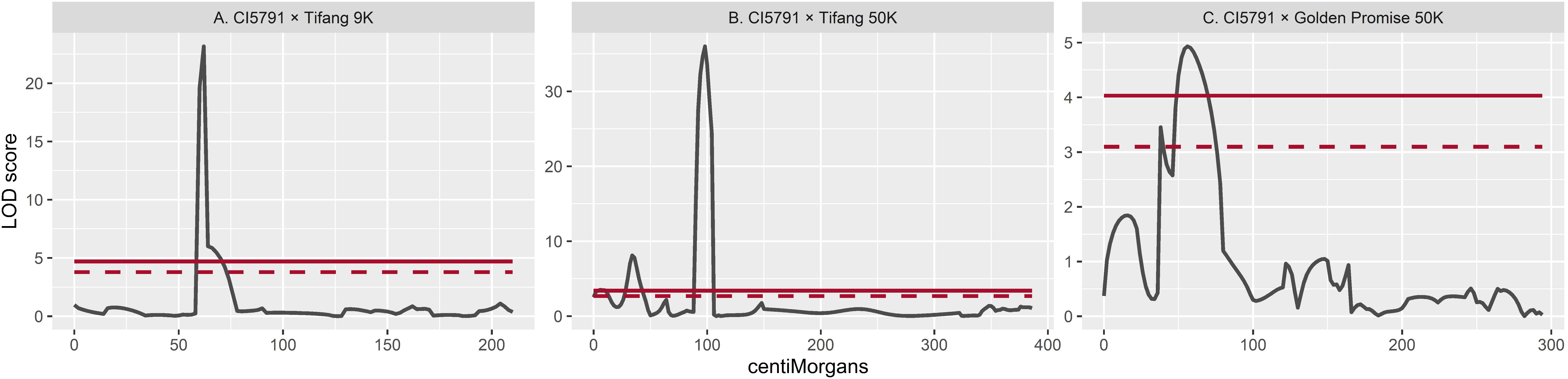
*Spt2* quantitative trait loci plots. for **A.** CI5791 × Tifang using 9K array derived markers; **B.** 50K array derived markers; and **C.** CI5791 × Golden Promise using 50K array derived markers along chromosome 5H. Black lines indicated LOD scores (y axis) along chromosome 5H of barley using centriMorgans. Dashed and solid red lines indicate 0.05 and 0.01 significant thresholds.

**Figure 3.**
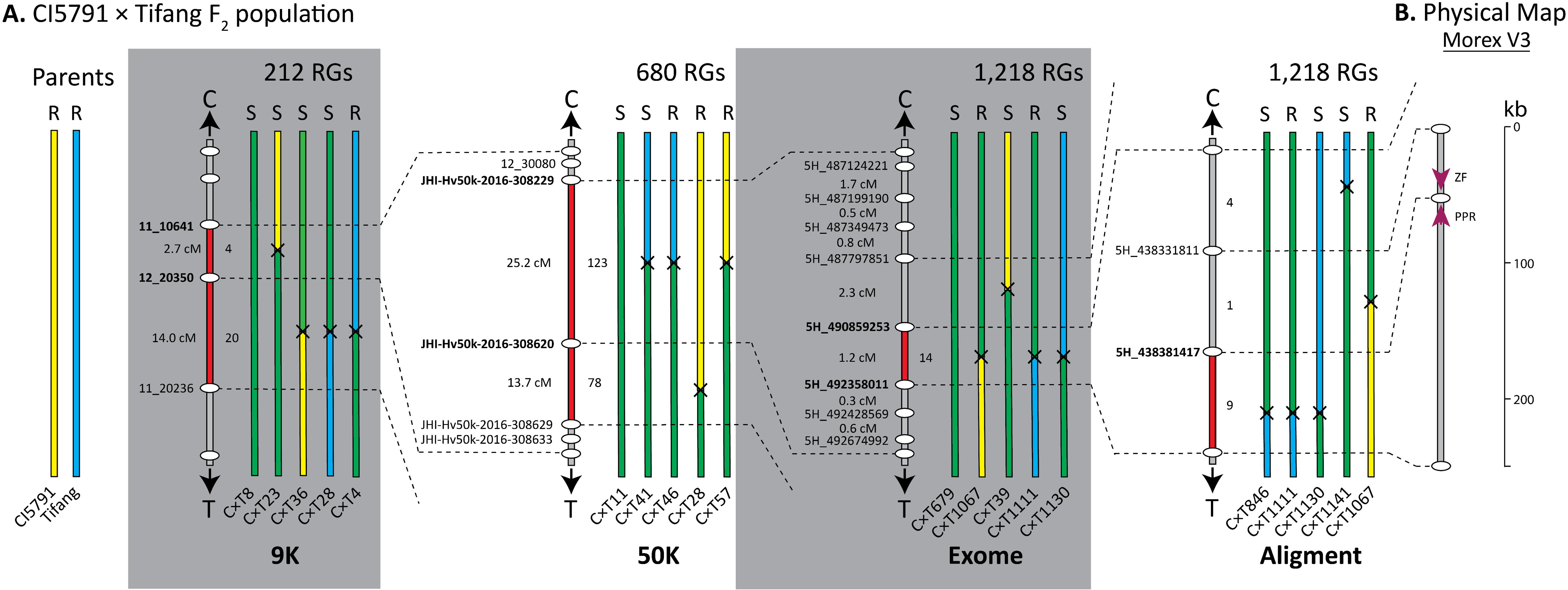
Genetic mapping of the *Spt2* locus within the CI5791 × Tifang population. **A.** Sequential mapping utilizing increased marker saturation and number of recombinant gametes corresponding to 9K, 50K, and exome capture sequencing and whole genome alignment derived marker panels. A representative set of F_2_ progeny are shown at each mapping stage with the specific accession ID indicated below the chromosome, whether the accession was resistant (R) or susceptible (S) and the total number of recombinant gametes (RGs) screened are indicated above. The parental sequences are shown in yellow (CI5791) or blue (Tifang), and heterozygous sequences in green. Black crosses indicate that recombination occurred between the two markers but do not indicate approximate location. The delimited region is marked as red along the chromosome and marker names in bold. Note that progeny numbers were reset from the second round of mapping. **B.** Physical interval within Morex V3 with genes shown as magenta arrows.

### High-Resolution CI5791 × Tifang Genetic Mapping with SNPs Derived from the 50K SNP Array

Within the genomic interval initially delimited via genetic mapping a total of 22 new SNP markers were identified as polymorphic between CI5791 and Tifang using the 50k iSelect genotyping platform. The first round of high-resolution mapping utilized a total of 340 CI5791 × Tifang F_2_ individuals (680 recombinant gametes) that were further saturated by successful amplification of an additional 20 PCR-GBS SNP markers from the 22 SNPs identified as polymorphic between the two parental lines. QTL analysis detected a highly significant locus on chromosome 5H between markers 12_30080 and JHI-Hv50k-2016-308620 with the most significant marker JHI-Hv50k-2016-308229 giving a LOD score of 36.0 (**Figure 2B**). LOD thresholds were calculated at 2.63 (α = 0.05) and 3.401 (α = 0.01). The 38.9 cM genetic interval (**Figure 3**) corresponds to a 5.1 Mb genomic interval within Morex V3 reference genome assembly with 64 annotated genes [27].

### High-Resolution CI5791 × Tifang Genetic Mapping with SNPs Derived from Exome Capture

The next round of high-resolution mapping utilized 609 individuals and 8 additional PCR-GBS markers derived from exome capture sequencing to saturate the genomic interval identified from 50K derived high-resolution mapping. One major QTL was detected within the chromosome 5H genomic interval between markers 5H_490859253 and 5H_492358011 with a peak LOD score of 85.4. LOD thresholds were calculated at 1.599 (α = 0.05) and 2.399 (α = 0.01). The 1.2 cM genetic interval (**Figure 3**) corresponds to a 1.5 Mb genomic interval within the Morex V3 reference genome assembly with nine annotated genes [27].

### Initial CI5791 × Golden Promise Genetic Mapping with Genotyping-by-Multiplex Sequencing

The CI5791 × Golden Promise population consisted of 83 F_2_ individuals and 180 markers spread across the barley genome, with 23 anchored to chromosome 5H of the Morex V3 reference genome [27]. Within this population, only one QTL was detected on chromosome 5H between markers 12_21036 and JHI-Hv50k-1016-311034 with a LOD score of 4.93 (**Figure 2C**). LOD thresholds were calculated at 3.098 (α = 0.05) and 4.031 (α = 0.01). This 31 cM genetic interval corresponds to a 33.3 Mb physical interval, encompassing the previously delimited locus identified within the CI5791 × Tifang population (**Figure 4**).

**Figure 4.**
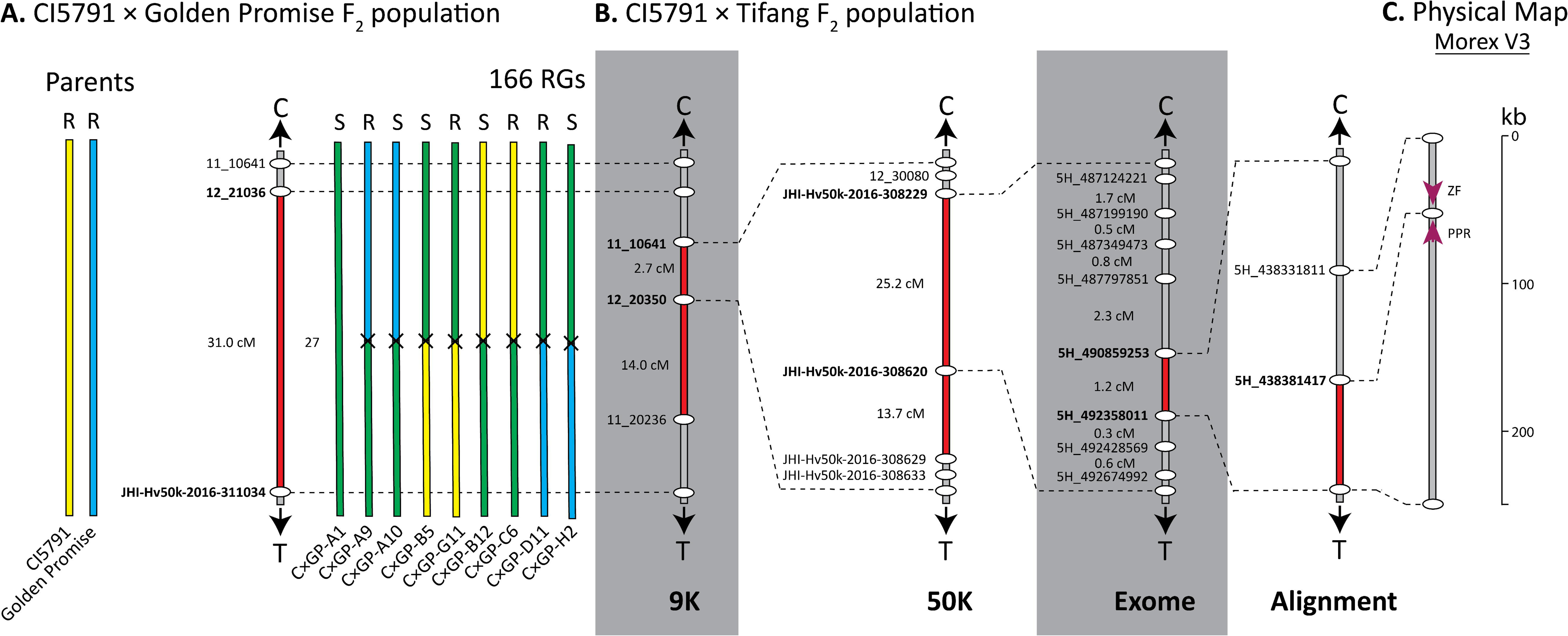
Genetic mapping of the *Spt2* locus within the CI5791 × Golden Promise population. **A.** A single F_2_ population using a 50K derived marker panel in comparison to the CI5791 × Tifang *Spt2* mapping population. A representative set of F_2_ progeny are shown at each mapping stage with the specific accession ID indicated below the chromosomes, whether the accession was resistant (R) or susceptible (S) and the total number of recombinant gametes (RGs) are indicated above. The parental sequences are shown in yellow (CI5791) or blue (Golden Promise), and heterozygous sequences in green. Black crosses indicate that recombination occurred between the two markers but do not indicate approximate location. The delimited region is marked as red along the chromosome and marker names in bold. **B.** The refined mapping of the *Spt2* locus using the CI5791 × Tifang F_2_ mapping population. **C.** Physical interval within Morex V3 with genes shown as magenta arrows.

### CI5791 Genome Assembly and *Spt2* Refinement

The CI5791 genome assembly was sequenced to a depth of 50x. Reads were assembled into 23,457 contigs (N50 of 309 kb) and scaffolded into the 12 pseudomolecule chromosomes of the Morex V3 genome assembly. Alignment of the *Spt2* region from CI5791 and Golden Promise identified multiple polymorphism that could be targeted with PACE. A total of two from 41 polymorphisms targeted were successfully converted to PACE and polymorphic between CI5791 and Tifang for analysis of the remaining 14 critical recombinants. From the four susceptible critical recombinants, one delimits the *Spt2* region to a single candidate gene annotated in the Morex V3 assembly as a PPR. A further four of the ten resistant critical recombinants also delimit the *Spt2* region to the PPR gene. Therefore, the *Spt2* interval was delimited between the markers 5H_438381417 and 5H_492358011 (∼198 kb) by five critical recombinants (**Figure 3**).

### Pangenome Assessment

The genomic interval delimiting *Spt2* between markers 5H_438381417 and 5H_492358011 corresponds to a ∼198 kb physical interval within the Morex V3 assembly and ranged from 177 to 266 kb within all the currently released barley genomes (**Table 1**). The parental lines CI5791 and Golden Promise harbor ∼217 and ∼200 kb physical intervals, respectively. Whereas the parental line Tifang does not currently have sequence data available. The CI5791 *Spt2* region is one the longest identified to date and the Golden Promise interval remains in concordance with other Western cultivars (**Table 1**). Both CI5791 and Golden Promise share high collinearity and synteny with Morex with only a deletion detected in the third syntenic block of CI5791 and a small inversion detected within Golden Promise when compared to CI5791 and Morex (**Figure 5A**). With the except of Haruna Nijo, a Japanese malting line, phylogenetic analysis of the *Spt2* locus reveals two main monophyletic groups separating European accessions and one American accession Morex with Asian and African accessions (**Figure 5B**). Canadian accession AAC Synergy and American accession Hockett form their own clades.

**Figure 5.**
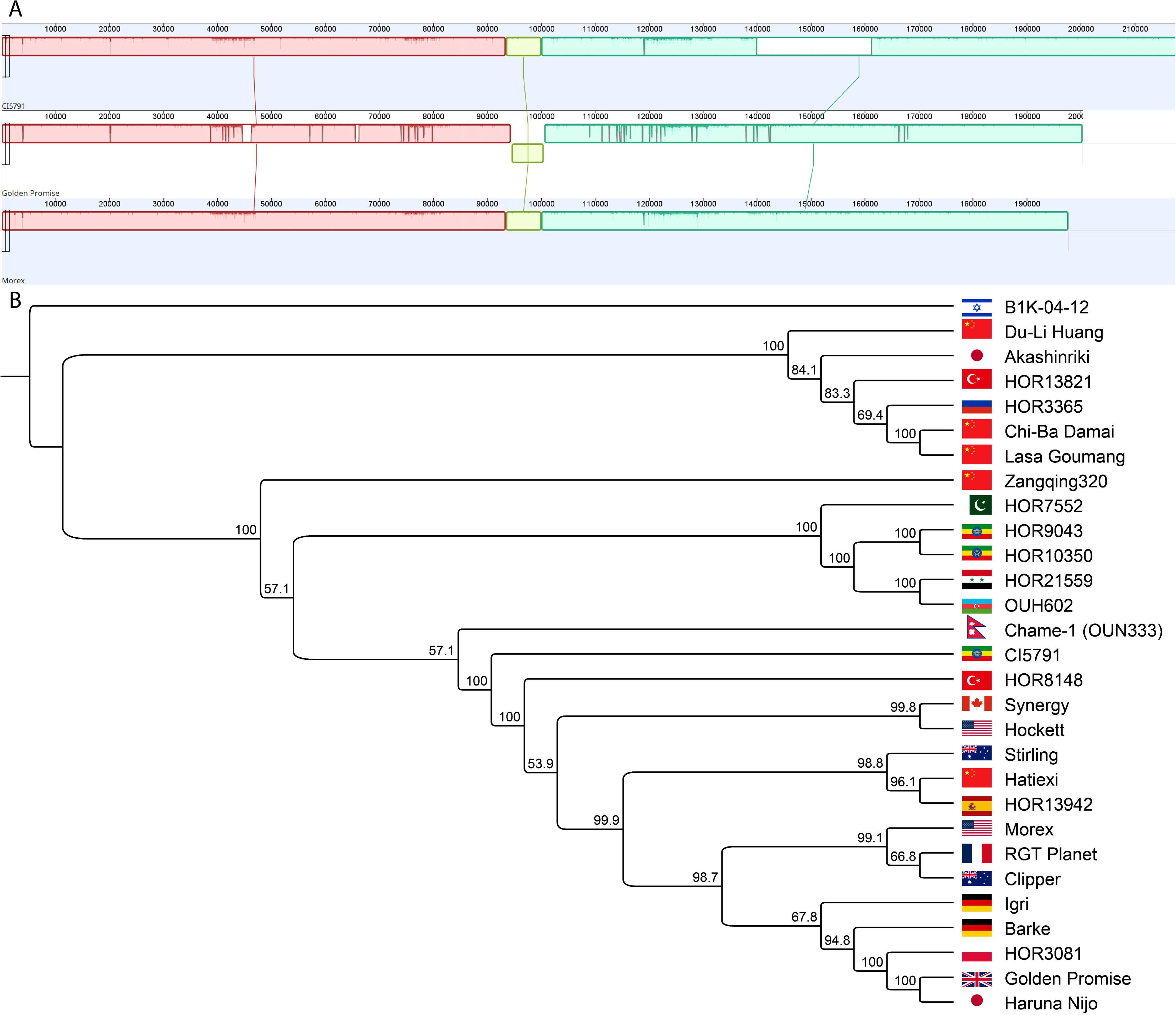
Pangenome comparisons of the *Spt2* region. **A.** Mauve alignment of the *Spt2* locus in CI5791 (top) and Golden Promise (middle), against the reference genome assembly of Morex V3 (bottom). Different colors indicate different syntenic colinear blocks. Blocks above and below the central line indicate sense and antisense. **B.** Phylogenetic tree using the *Spt2* locus from all 25 released barley genomes using MAFFT for alignment and RAxML for tree construction.

**Table 1.**
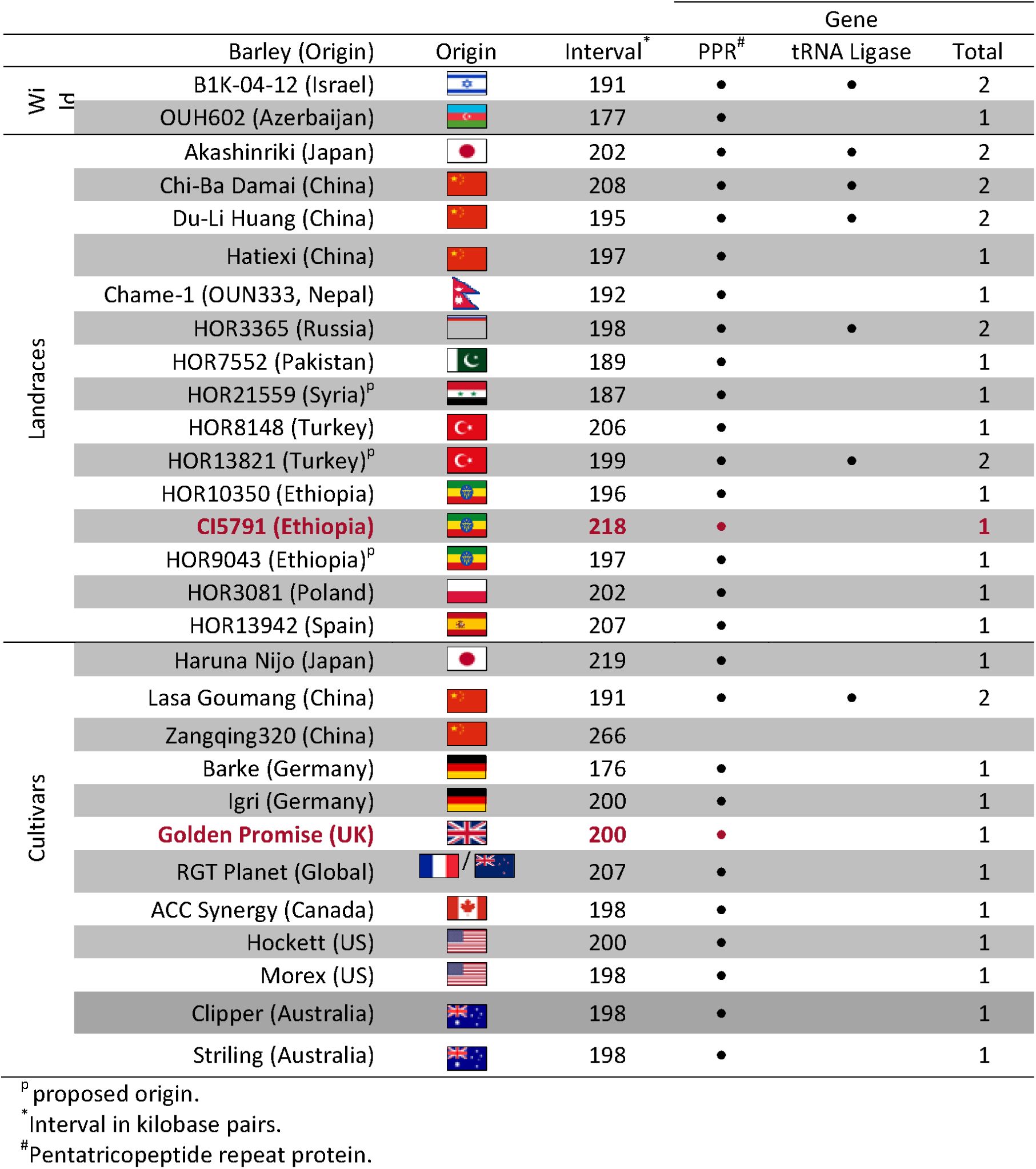
Annotated genes within the CI5791 genome assembly and all currently released pseudochromosome level barley genomes, organized by wild, landrace and cultivated varieties and country of origin. Red (parental line) and gray (other) dots indicate that the gene is present within the genome.

There are between one and two genes present within individual barley accessions, with a total of two unique genes present collectively across the delimited *Spt2* locus in all currently released pseudochromosome level genomes (**Table 1**). These genes include the PPR previously annotated in the Morex V3 assembly and a lysine tRNA ligase. However, only the PPR is consistently annotated in the majority of barley lines and the remaining lysine tRNA ligase gene is present in only seven wild and landrace accessions. The PPR is present in all lines except cultivar Zangqing320 from China. When comparing the PPR between CI5791 and Golden Promise, neither gene contains exonic polymorphisms, and therefore are predicted to have conserved amino acid sequences of the translated protein (**Table 2**). The PPR contains polymorphism within the intronic region, however these are not predicted to alter the splice site junctions (**Table 2**). Lastly, the PPR contains polymorphism upstream and downstream of the gene that may affect gene expression. The lysine tRNA ligase is not present within the *Spt2* delimited region in either CI5791 or Golden Promise.

**Table 2.** Polymorphisms identified within the *Spt2* candidate gene and respective consequence on gene structure between CI5791 and Golden Promise. Promotor and enhancer regions are covered by assessing homology 2kb up and downstream of the annotated gene unless otherwise stated.

| Gene | Exon | Intron | Downstream | Upstream | Consequence |
| --- | --- | --- | --- | --- | --- |
| Pentatricopeptide Repeat Protein | - | 2x SNP | 1x SNP | 2x SNP (2.4kb) | Unknown |
|  |  | 8x 1bp indel | 3x 1bp indel | 1x 1 bp indel |  |
|  |  | 101bp indel |  |  |  |

## Discussion

The *Spt2* gene which confers hybrid susceptibility was mapped to a single locus, whereby individuals heterozygous at the *Spt2* locus exhibit significantly increased susceptibility compared to either parent that are both resistant to the *Ptm* isolate 13IM8.3. This is the first documented case of non-transgressive segregation that supports the hybrid susceptibility model where susceptibility only manifests in the heterozygous state [28]. The *Spt2* locus is therefore non-transgressive as the locus was not mapped in previous mapping studies that utilized the CI5791 × Tifang recombinant inbred line (RIL) population and the same *Ptm* isolate 13IM8.3 [29]. In addition, this phenomenon is distinct from post-zygotic genetic incompatibilities such as hybrid necrosis that involves epistatic interactions between at least two loci and occur despite the absence of pathogen [25, 26], as no hybrid necrosis was observed during the development of the CI5791 × Tifang RIL population [30].

The *Spt2* locus was mapped to the same region within two independent populations that share CI5791 as a common parent. This raises the question of whether the same phenomena would exist in a Tifang × Golden Promise population and whether CI5791 is the main driver behind hybrid susceptibility. Due to the marker pool used on the CI5791 × Golden Promise population, the *Spt2* locus encompasses a 33.3 Mb physical region, delimited by the flanking markers 12_21036 and JHI-Hv50k-2016-311034 located at 428 MB and 461 Mb, respectively within the Morex V3 genome assembly [27]. Both 12_21036 and JHI-Hv50k-2016-311034 are the closest markers within the marker pool to the CI5791 × Tifang *Spt2* locus and could therefore not be refined further. In the CI5791 × Tifang population, the *Spt2* locus was sequentially mapped to 23.1 Mb (430.6 – 453.7 Mb), 5.1 Mb (434.0 – 439.1 Mb), 1.5 Mb (437.0 – 438.6 Mb), and ultimately a 198 kb (438.4 – 438.6 Mb) interval. The *Spt2* interval encompasses a single high confidence gene predicted to encode a PPR present in the publicly available Golden Promise [31], and Morex V3 [27] genome assemblies, and the CI5791 genome assembly generated for this study. As *Spt2* mapped to the same region in the CI5791 × Tifang and CI5791 × Golden Promise populations and comparative analysis of the CI5791 and Golden Promise genome sequences identified highly colinear, syntenic regions, we are confident this region represents *Spt2* in both populations.

Based on function and the current body of literature, the PPR candidate gene could be implicated in the heterozygous susceptibility phenomenon. Loss-of-function and knockdown mutants of PPR proteins in *Arabidopsis* have previously been implicated in increased susceptibility to bacterial and fungal pathogens [32]. PPR proteins are known to be found at exceptionally higher levels in plant organelles and shown to bind single stranded RNA in a similar manner to transcription activation-like effectors to DNA [33]. These PPR proteins are hypothesized to be at increased levels in plant organelles due to the lack of promoters in organelle genomes and therefore rely on PPR proteins to regulate gene expression at the RNA level affecting transcription, processing, splicing, stability, editing and translation

Heterosis has typically been explained by classical genetic models such as dominance, overdominance and epistasis of multiple genes [34]. As *Spt2* has been mapped to a single locus and comparative analysis determined that the PPR does not contain polymorphisms within exons that will change the primary amino acid sequence of the proteins, we therefore hypothesize that the causal gene underlying the locus results in hybrid susceptibility is due to insufficient expression, or overexpression of encoded protein/s caused by the polymorphisms in regulatory components. The PPR gene contains polymorphisms upstream, downstream and within introns between CI5791 and Golden Promise that may therefore affect the transcriptional regulation of the heterozygous alleles [35, 36]. These upstream, downstream and/or intronic polymorphisms could explain differential expression of the *Spt2* gene when present in a heterozygous state. We do not currently hypothesize that post-transcriptional modification of the proteins or alterations of protein complexes are affected as and the PPR is predicted to contain identical primary amino acid sequence in both the CI5791 and Golden Promise parents.

In addition, heterosis has often been associated with the genetic distance of the two parents, in that interspecies crosses exhibit increased heterosis over intraspecies crosses [34]. However, there is a growing body of evidence showing that this does not hold true and heterosis can be explained by epigenetic factors in intraspecies crosses [37–39]. For example, reports in *Arabidopsis* have shown that despite near identical genomes, different small interfering RNAs (siRNAs) and methylation states of each parent can produce F_1_ progeny with approximately 250% increased biomass [37]. Therefore, due to the fact that a single locus exhibiting limited genetic polymorphism within the PPR gene has been identified within an intraspecies cross, we also hypothesize that hybrid susceptibility could be the result of altered gene expression caused by altered siRNA, methylation states or histone modification at the *Spt2* locus within heterozygous progeny. Subsequently, the identified genetic polymorphisms at the *Spt2* locus are merely linked to these altered states when in the heterozygous state and not causal to the *Spt2-* mediated susceptibility. This altered expression of the *Spt2* gene in the heterozygous state, either suppressed or increased expression could result in loss of a resistance factor or the overexpression of a susceptibility target, respectively.

Interestingly, the *Spt2* locus colocalizes with the previously mapped loci *NBP_QRptt5-1* [40], *QRpt-5H.2* [41], and potentially an unnamed QTL [16, 42]. Therefore, we propose the primary locus designation as *Resistance to Pyrenophora teres 10* (*Rpt10*), with the secondary locus designation as *Spt2* (*Susceptibility to Pyrenophora teres 2*). Thus, *Rpt10/Spt2* may represent a resistance locus when in a homozygous state and a susceptibility locus when held in specific heterozygous states. However, further genetic analysis and gene characterization would need to be conducted to verify if the *Spt2* locus definitively colocalizes with the *Rpt10* locus and represents a singular *Rpt10/Spt2* locus similar to the *Rpt5/Spt1* locus in the barley-*Ptt* pathosystem [15].

With farmers increasingly relying on hybrid barley to maximize agronomic potential, discovery of the mechanisms underlying hybrid susceptibility is paramount to avoid the inadvertent release of hybrids with deleterious phenotypes. Hybrid barley is seen as a potential solution to combat stagnating yield increases, particularly in European feed lines, but could be applied to malting barley in the face of climate change [1, 3]. To our knowledge, this is the first documented case of non-transgressive, hybrid susceptibility within a single plant species, which could have grave consequences for hybrid cultivars if not addressed prior to release. Additional questions arise from the mechanism by which this hybrid susceptibility functions and whether genetic or epigenetic polymorphisms drive this mechanism. Elucidating this hybrid susceptibility mechanism will allow the identification of suitable and unsuitable parents in hybrid barley breeding programs and the release of both high quality and resistant hybrids.

## Conclusion

A rare non-transgressive hybrid susceptibility locus designated *Spt2* was high resolution mapped to a single candidate gene on chromosome 5H of barley using multiple rounds of mapping in a CI5791 × Tifang population. This non-transgressive hybrid susceptibility was also identified in a second population with Golden Promise sharing CI5791 as a common parent to the same locus. This single candidate is predicted to encode a pentatricopeptide repeat containing protein that has been previously implicated in susceptibility in *Arabidopsis* to bacterial and fungal pathogens. The *Spt2* locus was previously mapped as a resistance locus and therefore may function under a resistance mechanism when held in a homozygous state and a susceptibility locus when held in the heterozygous state. This study provides a base to further characterize and validate the *Spt2* locus and provides additional knowledge to understand the molecular mechanisms underlying hybrid susceptibility and vigor.

## Materials and Methods

### Biological Materials and Phenotyping

To reduce environmental variability all barley seedlings were grown in Argu soil in environmentally controlled growth chambers (Conviron, Winnipeg, Canada) under a 12 hr light/12 hr dark cycle, at 21°C. The F_2_ seedlings used for genetic mapping of *Spt2* were derived from CI5791 × Tifang or CI5791 × Golden Promise crosses. CI5791 is a 2-row Ethiopian landrace, Tifang is a 6-row Chinese landrace, and Golden Promise is a Scottish 2-row malting cultivar. The F_1_ individuals were self-fertilized to generate F_2_ seed and individual F_2_ seed were planted in cone-tainers for the first two rounds of mapping, and 96-cell trays for the final round of mapping using the CI5791 × Tifang population and the CI5791 × Golden Promise population. Phenotyping could not be repeated due to the genetic nature of the *Spt2-*mediated hybrid susceptibility which required mapping with F_2_ individuals. Successive steps of higher resolution mapping and increased marker saturation were performed utilizing additional F_2_ progeny from the CI5791 × Tifang cross. For initial mapping 106 CI5791 × Tifang F_2_ individuals (representing 212 recombinant gametes) were phenotyped and genotyped with SNP markers derived from the 9K Illumina SNP data [43]. This was followed by higher resolution genetic mapping with 340 F_2_ individuals (680 recombinant gametes) that were genotyped with SNP markers generated utilizing the 50K Illumina SNP data [44]. The next round of high-resolution mapping utilized 609 F_2_ individuals (1,218 recombinant gametes) and further saturation of markers utilizing exome capture data generated on the critical recombinant individuals identified with the 50K Illumina SNP data [44, 45]. The last step in high-resolution mapping utilized 14 critical recombinants identified in the previous screening of 609 CI5791 × Tifang F_2_ individuals saturated with additional markers identified as polymorphic from the CI5791 and Golden Promise [31, 46] genomic alignment. Each step of genetic mapping was conducted independently with exception of the last rounds within the CI5791 × Tifang high-resolution population which utilized DNA from individuals identified in the previous screen as well as the same phenotypic data for further marker saturation in the regions identified with critical recombination events. A total of 83 F_2_ individuals were used for the CI5791 × Golden Promise F_2_ population.

One-week post planting of each of the F_2_ populations, dried agar plugs of the *Ptm* isolate 13IM.3 were placed on plates of V3 media and allowed to grow for approximately seven days in the dark before exposing them to a 24 hr light/24 hr dark cycle to induce sporulation. The *Ptm* isolate 13IM.3 was collected from Blackfoot, Idaho in 2013 [47]. After sporulation was induced the plates were flooded with ∼10 ml of ddH_2_0 and scraped using a sterile inoculating loop. The conidia spores collected in the water suspension were counted under a microscope using a 5 μl droplet and adjusted to ∼2,000 spores/ml with one drop (∼30 μl) of Tween 20 surfactant per 50ml. Plants were spray inoculated at approximately 20 psi until runoff at the second leaf stage which was ∼14 days post planting under growth chamber conditions, placed in a mist chamber with 24 hr light conditions and misted to maintain 100% relative humidity. After 24 hrs in the mist chamber, the inoculated seedlings were allowed to air dry and moved back to the growth chamber set at a 12 hr light/12 hr dark cycle, and 21°C. Seven days post inoculation plants were phenotypically scored using the 1-5 SFNB rating scale described by [48]. Individuals were considered resistant when less than 3 on the rating scale and susceptible when 3 or above on the rating scale.

### DNA Extraction for Genotyping

Tissue from individual seedlings were collected in 96-deep well plates prior to inoculation and DNA extracted for 9K derived initial and 50K derived high-resolution mapping using PCR genotyping-by-sequencing (PCR-GBS) as described by [15] and [49] using the Ion Torrent PGM platform. For the final round of high-resolution mapping and low-resolution mapping with the CI5791 × Golden Promise population, the leaf tissue was collected in sample plates, frozen at −80°C, lyophilized and DNA extracted using the oKtopure™ automated DNA extraction platform following the manufacturer’s instructions.

### CI5791 × Tifang 9K Derived Initial Mapping

For initial genetic mapping using CI5791 × Tifang F_2_ individuals a custom panel of 365 SNPs (**Supplemental File 1**) was utilized. The SNP markers utilized to develop the custom PCR-GBS panel for amplicon sequencing were derived from the barley 9k iSelect SNP platform data [43]. The 365 SNPs were selected based on polymorphism between CI5791 and Tifang as well as uniform distribution across the barley genome. Primers were designed from the source sequences extracted from the T3/Barley database (https://barley.triticeaetoolbox.org) to produce ∼125 bp amplicons over the targeted SNPs. Primers were tagged with CS1 and CS2 adapters (Fluidigm, San Francisco CA) for forward and reverse primers, respectively. Each primer was tested individually for amplification and primers pooled to generate ten multiplex pools. Subsequently, PCR-GBS was performed as described by [15] and [49] on the Ion Torrent PGM platform. Reads were trimmed by 22nt in CLC Genomics Workbench to remove adapters. Trimmed reads were aligned to the reference amplicon panel using bwa-mem [50], converted and sorted to BAM files using samtools [51] and SNPs called using the Genome Analysis Toolkit HaplotypeCaller tool [52, 53]. SNPs were filtered using vcftools [54] using the following filters *--minQ 30 --minDP 3* and *--max-missing 0.5*. MapDisto 2.1.7 [55] was used for genetic map construction and file export for QTL mapping in QGene 4.4.0 [56] using the composite interval mapping algorithm and default parameters for selecting cofactors (**Supplemental File 2**). LOD thresholds at α level = 0.05 and α level = 0.01 were calculated using a 1000 permutation test. LOD scores and significance thresholds were parsed for visualization using ggplot2 within *R* 4.1.2.

### CI5791 × Tifang 50K Derived High-Resolution Mapping

Following initial mapping, both parental lines were subjected to the barley 50k iSelect genotyping platform [44]. Molecular markers within the region delimited by the initial genetic mapping were analyzed for polymorphisms between CI5791 and Tifang. The analysis identified 22 polymorphic SNPs (**Supplemental File 3**) based extracted from the T3/Barley database (https://barley.triticeaetoolbox.org). Primers for the RAD-GBS panel were designed using Primer3-Plus within Geneious Prime® (https://www.geneious.com), and checked for specificity using BLAST on the EnsemblPlants [57] using the barley genome. Multiplex robustness was optimized by detecting predicted primer dimer and multiplex ability using MFE-Primer [58]. The PCR-GBS and SNP calling pipeline followed the same protocol as the initial mapping except reads were trimmed using Trimmomatic [59]. High-resolution QTL mapping was performed in the same manner as the initial mapping described above (**Supplemental File 4**).

### CI5791 × Tifang Exome Capture Derived High-Resolution Mapping

Five lines exhibiting recombination within the delimited region were selected for exome capture. SeqCap EZ HyperCap exome capture [45] was performed as per the manufacturer’s instructions (Roche Sequencing Solutions, Pleasanton, CA, USA) with the following adjustments. Starting DNA concentration was increased from the recommended 100ng to 300ng per sample based on Hoescht DNA quantification and pooled DNA concentration was increased from 1 µg to 2µg. The final purified pool was sequenced on a single flow cell on the Illumina NextSeq®500 sequencing platform. A total of 46 polymorphic SNPs (**Supplemental File 5**) were identified from the exome capture sequencing data and oligonucleotide primer pairs were designed as previously described for the 9K SNP source sequences with the following adjustments. The 5’ termini of the primers were tagged with M13 and P1/B sequences to facilitate barcoding. DNA from the 340 F_2_ individuals from the 50K derived high-resolution mapping and an additional 269 F_2_ individuals were utilized for a total of 609 F_2_ individuals representing 1,218 recombinant gametes. Sequencing was performed using an Ion PI™ Chip on the Ion Proton™ system platform. Individual fastq files were parsed using fastq-grep command due to the unequal barcode lengths used to increase the number of samples in the multiplex sequencing library. SNP calling preceded as described in 50K derived high-resolution mapping and QTL mapping was performed in the same manner as 9K derived initial mapping as described above (**Supplemental File 6**).

### CI5791 × Golden Promise Genotype-by-Multiplex Sequencing Mapping

A total of 83 CI5791 × Golden Promise F_2_ individuals were screened for disease response as described in biological materials and phenotyping. After DNA extraction, accessions were genotyped using a genotype-by-multiplex sequencing marker panel [60] using Illumina sequencing. SNP calling preceded as described in exome derived high-resolution mapping of CI5791 × Tifang except the reads were trimmed using fastp [61]. QTL mapping was performed in the same manner as 9K derived initial mapping as described above (**Supplemental File 7**).

### CI5791 Genome Assembly

The barley accession CI5791 was subjected to whole genome sequencing for genome assembly. Briefly, barley was grown to the three-leaf stage and dark treated for 24 hr. Leaves were cut, and 50 g of young leaf tissue was weighed and wrapped in aluminum foil and immediately submerged in liquid nitrogen. Tissue was subsequently stored at −80°C. High molecular weight DNA was extracted, and sequencing performed using one SMRT Cell 8M in Continuous Long Read mode of the PacBio Sequel II System at the University of Arizona Genomics Institute. Raw reads were assembled into contigs using Canu 2.2.1 [62], and scaffolded to the Morex V3 reference genome assembly [27].

### *Spt2* Refinement using CI5791 and Golden Promise Polymorphisms

Fourteen critical recombinants were identified from the 609 CI5791 × Tifang F_2_ individuals screened within the delimited 1.2 cM (∼1.5 Mb) *Spt2* region that could be utilized for additional marker saturation based on identified polymorphisms. As all identified polymorphisms were exhausted from previous genotyping platforms, the CI5791 genome assembly and the publicly available Golden Promise genome assembly were utilized to identify novel polymorphism to further saturate the region with genetic markers. To identify polymorphisms between CI5791 and Golden Promise, 500 kb intervals of the *Spt2* locus were aligned using MAFFT [63]. SNPs and 1 bp indels between CI5791 and Golden Promise were targeted for PACE (3CR Bioscience, Harlow, UK) primer design within Geneious Prime® 2023.2.1 (https://www.geneious.com). Two polymorphisms from the 41 identified were targeted and successfully converted to PACE genotyping assays (**Supplemental File 8**). Primer designs followed 3CR Bioscience guidance, while minimizing Tm primer differences to less than 1 **°**C and avoiding homopolymer runs. Primer specificity was determined using EnsemblPlants [57] BLAST tool and the Morex V3 genome assembly. Oligonucleotides were synthesized (Sigma-Alrich, St. Loius, MO, USA) and diluted to 100 μM stocks. Primer concentrations followed manufacturers recommendations with working primer assay stocks created by combining 12 µl of each forward primer, 30 µl of the common reverse primer, and 46 µl of water. Assay mastermixes were produced by combining a total of 5.0 μl standard ROX PACE mastermix (3CR Bioscience, Harlow, UK) and 0.14 µl of working primer assay per sample. Cycling conditions were optimized by performing 10 µl reactions containing 5.0 µl assay mastermix and 5.0 µl of parental DNA and two non-template water controls (NTCs) in 384-well plate format on a CFX384 qPCR thermocycler (Bio-Rad Laboratories, Hercules, CA, USA). Cycle number was optimized by assessing every two cycles between 26-36x cycles post touchdown of 10 cycles (61–55°C, −0.6°C/cycle). Assays were deemed successful if two distinct homozygous clusters were obtained with one group consisting of CI5791 samples and the second group consisting of Golden Promise and Tifang samples. The successfully converted assays were subsequently performed on the critical recombinants using identical conditional with two samples of each parent, two CI5691 × Tifang F_1_ samples and two NTC samples as controls. All results were analyzed in CFX Maestro 1.1 (Bio-Rad Laboratories) (**Supplemental File 9**).

### Pangenome Assessment

Following the *Spt2* region delimitation utilizing polymorphisms identified from CI5791 and Golden Promise genome alignment, flanking regions of the *Spt2* interval of Morex V3 [27] were used as seeds to BLAST against each barley genome within the barley pangenome [46], additionally released pseudochromosome level genomes of cultivars Golden Promise [31, 46], AAC Synergy [64], Clipper, Stirling [65], Haruna Nijo [66], Lasa Goumang [67], Zangqing320 [68], wild barley OUH602 [69], landrace Hatiexi [70] and landrace CI5791 from this study. The *Spt2* regions from the sequences of all the barley lines were compared to assess the gene space diversity. Divergence of the *Spt2* locus between all 25 barley genome assemblies was constructed using MAFFT [63] and RAxML [71] within Geneious Prime® 2023.2.1 (https://www.geneious.com). Additionally, the *Spt2* region of Morex V3 [27], Golden Promise [31, 46] and CI5791 were compared using Mauve [72] using default settings within Geneious Prime® 2023.2.1.

## Availability of data and materials

The CI5791 genome assembly is publicly available under BioProject ID <u>PRJNA1088520</u>. The raw sequencing data generated for PCR-GBS and GMS genotyping is publicly available as Sequence Read Archives database under BioProject IDs <u>PRJNA1088606</u>, <u>PRJNA1088747</u>, <u>PRJNA1088752</u> and <u>PRJNA1088849</u>. The exome capture sequencing is publicly available as a Sequence Read Archive under BioProject ID <u>PRJNA1088748</u>.

## Abbreviations

HR: Hypersensitive Response
LOD: Logarithm of Odds
NE: Necrotrophic Effector
NFNB: Net Form Net Blotch
PACE®: PCR Allele Competitive Extension®
PCD: Programmed Cell Death
PCR-GBS: Polymerase Chain Reaction – Genotyping-by-Sequencing
*Ptm*: *Pyrenophora teres* f. *maculata*
*Ptt*: *Pyrenophora teres* f. *teres*
QTL: Quantitative Trait Loci
R Gene/Protein: Resistance Gene/Protein
RLK: Receptor-Like Kinase
RLP: Receptor-Like Protein
Rpt#: Resistance to *Pyrenophora teres* #
SFNB: Spot Form Net Blotch
SNP: Single Nucleotide Polymorphism
Spt#: Susceptibility to *Pyrenophora teres* #
V8-PDA: V8 Juice – Potato Dextrose Agar
WRKY: Tryptophan (W)-Arginine (R)-Lysine (K)-Tyrosine (Y)

## Acknowledgements

We thank the Western Regional Small Grains Genotyping Laboratory (Deven See) and the Laboratory of Biotechnology and Bioanalysis (Derek Pouchnik) at Washington State University, specifically K. Marlowe for exome capture and Illumina sequencing, Travis Ruff and Mark Wildung for Ion Torrent sequencing, and Marcus Hooker for bioinformatic support. We would also like to thank undergraduates Sarah Harkins and Hannah Peha for project assistance and Dave Kudrna at the Arizona Genomics Institute of the University of Arizona for DNA extraction and PacBio sequencing of CI5791.

## Author Information

### Author Contributions

**Shaun Clare:** Conceptualization, Methodology, Software, Validation, Formal Analysis, Investigation, Data Curation, Visualization, Writing - Original Draft. **Abdullah Alhasel** : Methodology, Validation, Formal Analysis, Investigation, Data Curation. **Mengyuan Li:** Methodology, Software, Validation, Formal Analysis, Investigation, Data Curation. **Karl Effertz:** Investigation. **Roshan Sharma Poudel:** Methodology, Software, Validation, Formal Analysis, Investigation, Data Curation, **Jianwei Zhang:** Resources, Supervision, Funding Acquisition. **Robert Brueggeman** : Conceptualization, Methodology, Resources, Supervision, Project Administration, Funding Acquisition. **All Authors:** Writing - Review & Editing.

### Corresponding Author

Correspondence to Robert S. Brueggeman.

## Ethics Declarations

### Ethics approval and consent to participate

Experimental research and field studies on plants, including the collection of plant material, was in accordance with relevant institutional, national, and international guidelines and legislation.

### Consent for publication

Not applicable.

## Supplemental Information

**Supplemental File 1.** Amplicons amplified in the 365 panel, derived from the barley 9K iSelect platform for initial mapping in the CI5791 × Tifang F population.

**Supplemental File 2.** Mapping data for the CI5791 × Tifang F population using the 9K derived PCR-Genotyping-By-Sequencing protocol.

**Supplemental File 3.** Amplicons amplified from the barley 50K iSelect platform in the high-resolution mapping in the CI5791 × Tifang F population.

**Supplemental File 4.** Mapping data for the CI5791 × Tifang F population using the 50K derived PCR-Genotyping-By-Sequencing protocol.

**Supplemental File 5.** Amplicons amplified from the barley exome capture in the high-resolution mapping in the CI5791 × Tifang F population.

**Supplemental File 6.** Mapping data for the CI5791 × Tifang F population using the exome capture derived PCR-Genotyping-By-Sequencing protocol.

**Supplemental File 7.** Mapping data for the CI5791 × Golden Promise F population using the exome capture derived Genotyping by Multiplex Sequencing protocol.

**Supplemental File 8.** Primers for PACE assays used for the high-resolution mapping in the CI5791 × Tifang F population.

**Supplemental File 9.** Mapping data for the CI5791 × Tifang F population using PCR Allele Competitive Extension® assays.

## References

1. Longin CFH, Mühleisen J, Maurer HP, Zhang H, Gowda M, Reif JC. Hybrid breeding in autogamous cereals. Theoretical and Applied Genetics. 2012;125:1087–96.

2. Mühleisen J, Maurer HP, Stiewe G, Bury P, Reif JC. Hybrid Breeding in Barley. Crop Science. 2013;53:819–24.

3. Xie W, Xiong W, Pan J, Ali T, Cui Q, Guan D, et al. Decreases in global beer supply due to extreme drought and heat. Nature Plants. 2018;4:964–73.

4. Dreiseitl A. A novel way to identify specific powdery mildew resistance genes in hybrid barley cultivars. Scientific Reports. 2020;10:18930.

5. Kourelis J, van der Hoorn RAL. Defended to the Nines: 25 Years of Resistance Gene Cloning Identifies Nine Mechanisms for R Protein Function. The Plant Cell. 2018;30:285 LP – 299.

6. Johal GS, Briggs SP. Reductase activity encoded by the HM1 disease resistance gene in maize. Science. 1992;258:985 LP – 987.

7. Cesari S, Bernoux M, Moncuquet P, Kroj T, Dodds PN. A novel conserved mechanism for plant NLR protein pairs: the “integrated decoy” hypothesis. Frontiers in Plant Science. 2014;5:606.

8. van der Hoorn RAL, Kamoun S. From Guard to Decoy: A New Model for Perception of Plant Pathogen Effectors. The Plant Cell. 2008;20:2009 LP – 2017.

9. Kolodziej MC, Singla J, Sánchez-Martín J, Zbinden H, Šimková H, Karafiátová M, et al. A membrane-bound ankyrin repeat protein confers race-specific leaf rust disease resistance in wheat. Nature Communications. 2021;12:956.

10. Wang H, Zou S, Li Y, Lin F, Tang D. An ankyrin-repeat and WRKY-domain-containing immune receptor confers stripe rust resistance in wheat. Nature Communications. 2020;11:1353.

11. Tamang P, Richards JK, Solanki S, Ameen G, Sharma Poudel R, Deka P, et al. The Barley *HvWRKY6* Transcription Factor Is Required for Resistance Against *Pyrenophora teres* f. *teres*. Frontiers in Genetics. 2021;11:1762.

12. Liu Z, Holmes DJ, Faris JD, Chao S, Brueggeman RS, Edwards MC, et al. Necrotrophic effector-triggered susceptibility (NETS) underlies the barley–*Pyrenophora teres* f. *teres* interaction specific to chromosome 6H. Molecular Plant Pathology. 2015;16:188–200.

13. Abu Qamar M, Liu ZH, Faris JD, Chao S, Edwards MC, Lai Z, et al. A region of barley chromosome 6H harbors multiple major genes associated with net type net blotch resistance. Theoretical and Applied Genetics. 2008;117:1261.

14. Shjerve RA, Faris JD, Brueggeman RS, Yan C, Zhu Y, Koladia V, et al. Evaluation of a *Pyrenophora teres* f. *teres* mapping population reveals multiple independent interactions with a region of barley chromosome 6H. Fungal Genetics and Biology. 2014;70:104–12.

15. Richards J, Chao S, Friesen T, Brueggeman R. Fine mapping of the barley chromosome 6H net form net blotch susceptibility locus. G3 Genes|Genomes|Genetics. 2016;6:1809–18.

16. Clare SJ, Wyatt NA, Brueggeman RS, Friesen TL. Research advances in the *Pyrenophora teres*–barley interaction. Molecular Plant Pathology. 2020;21:272–88.

17. Richards J, Li J, Koladia V, Wyatt N, Rehman S, Brueggeman RS, et al. A Moroccan *Pyrenophora teres* f. *teres* population defeats *Rpt5*, the broadly effective resistance on barley chromosome 6H. Phytopathology®. 2024;114:193–9.

18. Liu Z, Ellwood SR, Oliver RP, Friesen TL. *Pyrenophora teres*: profile of an increasingly damaging barley pathogen. Molecular Plant Pathology. 2011;12:1–19.

19. Yuzon JD, Wyatt NA, Vasighzadeh A, Clare S, Navratil E, Friesen TL, et al. Hybrid inferiority and genetic incompatibilities drive divergence of fungal pathogens infecting the same host. Genetics. 2023;224:iyad037.

20. Steffenson B. Net blotch. In: Mathre D, editor. Compendium of Barley Diseases. The American Phytopathological Society: St. Paul Minnesota; 1997. p. 28–31.

21. Grewal TS, Rossnagel BG, Pozniak CJ, Scoles GJ. Mapping quantitative trait loci associated with barley net blotch resistance. Theoretical and Applied Genetics. 2008;116:529–39.

22. Murray GM, Brennan JP. Estimating disease losses to the Australian barley industry. Australasian Plant Pathology. 2010;39:85–96.

23. Uranga JP, Schierenbeck M, Perelló AE, Lohwasser U, Börner A, Simón MR. Localization of QTL for resistance to *Pyrenophora teres* f. *maculata*, a new wheat pathogen. Euphytica. 2020;216:1–13.

24. Koide Y, Sakaguchi S, Uchiyama T, Ota Y, Tezuka A, Nagano AJ, et al. Genetic Properties Responsible for the Transgressive Segregation of Days to Heading in Rice. G3 Genes|Genomes|Genetics. 2019;9:1655–62.

25. Bomblies K, Weigel D. Hybrid necrosis: autoimmunity as a potential gene-flow barrier in plant species. Nat Rev Genet. 2007;8:382–93.

26. Li L, Weigel D. One Hundred Years of Hybrid Necrosis: Hybrid Autoimmunity as a Window into the Mechanisms and Evolution of Plant–Pathogen Interactions. Annual Review of Phytopathology. 2021;59:213–37.

27. Mascher M, Wicker T, Jenkins J, Plott C, Lux T, Koh CS, et al. Long-read sequence assembly: a technical evaluation in barley. The Plant Cell. 2021;33:1888–906.

28. LeBoldus JM, Isabel N, Floate KD, Blenis P, Thomas BR. Testing the ‘Hybrid Susceptibility’ and ‘Phenological Sink’ Hypotheses Using the *P. balsamifera – P. deltoides* Hybrid Zone and Septoria Leaf Spot [*Septoria musiva*]. PLOS ONE. 2013;8:e84437.

29. Alhashel AF, Sharma Poudel R, Fiedler J, Carlson CH, Rasmussen J, Baldwin T, et al. Genetic mapping of host resistance to the *Pyrenophora teres* f. *maculata* isolate 13IM8.3. G3 Genes|Genomes|Genetics. 2021;11:jkab341.

30. Koladia VM, Faris JD, Richards JK, Brueggeman RS, Chao S, Friesen TL. Genetic analysis of net form net blotch resistance in barley lines CIho 5791 and Tifang against a global collection of *P. teres* f. *teres* isolates. Theoretical and Applied Genetics. 2017;130:163–73.

31. Schreiber M, Mascher M, Wright J, Padmarasu S, Himmelbach A, Heavens D, et al. A Genome Assembly of the Barley ‘Transformation Reference’ Cultivar Golden Promise. G3 Genes|Genomes|Genetics. 2020;10:1823–7.

32. Park YJ, Lee HJ, Kwak KJ, Lee K, Hong SW, Kang H. MicroRNA400-Guided Cleavage of Pentatricopeptide Repeat Protein mRNAs Renders *Arabidopsis thaliana* More Susceptible to Pathogenic Bacteria and Fungi. Plant and Cell Physiology. 2014;55:1660–8.

33. Yin P, Li Q, Yan C, Liu Y, Liu J, Yu F, et al. Structural basis for the modular recognition of single-stranded RNA by PPR proteins. Nature. 2013;504:168–71.

34. Lauss K, Wardenaar R, Oka R, van Hulten MHA, Guryev V, Keurentjes JJB, et al. Parental DNA Methylation States Are Associated with Heterosis in Epigenetic Hybrids. Plant Physiology. 2018;176:1627–45.

35. Haberle V, Stark A. Eukaryotic core promoters and the functional basis of transcription initiation. Nature Reviews Molecular Cell Biology. 2018;19:621–37.

36. Shaul O. How introns enhance gene expression. The International Journal of Biochemistry & Cell Biology. 2017;91:145–55.

37. Groszmann M, Greaves IK, Albertyn ZI, Scofield GN, Peacock WJ, Dennis ES. Changes in 24-nt siRNA levels in Arabidopsis hybrids suggest an epigenetic contribution to hybrid vigor. Proceedings of the National Academy of Sciences. 2011;108:2617–22.

38. Groszmann M, Greaves IK, Fujimoto R, James Peacock W, Dennis ES. The role of epigenetics in hybrid vigour. Trends in Genetics. 2013;29:684–90.

39. Springer NM. Epigenetics and crop improvement. Trends in Genetics. 2013;29:241–7.

40. Wonneberger R, Ficke A, Lillemo M. Identification of quantitative trait loci associated with resistance to net form net blotch in a collection of Nordic barley germplasm. Theoretical and Applied Genetics. 2017;130:2025–43.

41. Clare SJ, Çelik Oğuz A, Effertz K, Sharma Poudel R, See D, Karakaya A, et al. Genome-wide association mapping of *Pyrenophora teres* f. *maculata* and *Pyrenophora teres* f. *teres* resistance loci utilizing natural Turkish wild and landrace barley populations. G3 Genes|Genomes|Genetics. 2021;11.

42. Williams KJ, Platz GJ, Barr AR, Cheong J, Willsmore K, Cakir M, et al. A comparison of the genetics of seedling and adult plant resistance to the spot form of net blotch (*Pyrenophora teres* f. *maculata*). Australian Journal of Agricultural Research. 2003;54:1387–94.

43. Comadran J, Kilian B, Russell J, Ramsay L, Stein N, Ganal M, et al. Natural variation in a homolog of *Antirrhinum CENTRORADIALIS* contributed to spring growth habit and environmental adaptation in cultivated barley. Nature Genetics. 2012;44:1388.

44. Bayer MM, Rapazote-Flores P, Ganal M, Hedley PE, Macaulay M, Plieske J, et al. Development and Evaluation of a Barley 50k iSelect SNP Array. Frontiers in Plant Science. 2017;8:1792.

45. Mascher M, Richmond TA, Gerhardt DJ, Himmelbach A, Clissold L, Sampath D, et al. Barley whole exome capture: a tool for genomic research in the genus *Hordeum* and beyond. The Plant Journal. 2013;76:494–505.

46. Jayakodi M, Padmarasu S, Haberer G, Bonthala VS, Gundlach H, Monat C, et al. The barley pangenome reveals the hidden legacy of mutation breeding. Nature. 2020;588:284–9.

47. Clare SJ, Duellman KM, Richards JK, Sharma Poudel R, Merrick LF, Friesen TL, et al. Association Mapping Reveals a Reciprocal Virulence/Avirulence Locus Within Diverse US *Pyrenophora teres* f. *maculata* Isolates. BMC Genomics. 2022;23.

48. Neupane A, Tamang P, Brueggeman RS, Friesen TL. Evaluation of a barley core collection for spot form net blotch reaction reveals distinct genotype-specific pathogen virulence and host susceptibility. Phytopathology. 2015;105:509–17.

49. Sharma Poudel R, Al-Hashel AF, Gross T, Gross P, Brueggeman R. Pyramiding *rpg4*- and *Rpg1*-mediated stem rust resistance in barley requires the *Rrr1* gene for both to function. Department of Plant Pathology, North Dakota State University, Fargo, ND, United States.; 2018.

50. Li H, Durbin R. Fast and accurate short read alignment with Burrows–Wheeler transform. Bioinformatics. 2009;25:1754–60.

51. Danecek P, Bonfield JK, Liddle J, Marshall J, Ohan V, Pollard MO, et al. Twelve years of SAMtools and BCFtools. GigaScience. 2021;10:giab008.

52. McKenna A, Hanna M, Banks E, Sivachenko A, Cibulskis K, Kernytsky A, et al. The Genome Analysis Toolkit: A MapReduce framework for analyzing next-generation DNA sequencing data. Genome Research. 2010;20:1297–303.

53. Poplin R, Ruano-Rubio V, DePristo MA, Fennell TJ, Carneiro MO, Van der Auwera GA, et al. Scaling accurate genetic variant discovery to tens of thousands of samples. bioRxiv. 2018;:201178.

54. Danecek P, Auton A, Abecasis G, Albers CA, Banks E, DePristo MA, et al. The variant call format and VCFtools. Bioinformatics (Oxford, England). 2011;27:2156–8.

55. Heffelfinger C, Fragoso CA, Lorieux M. Constructing linkage maps in the genomics era with MapDisto 2.0. Bioinformatics. 2017;33:2224–5.

56. Joehanes R, Nelson JC. QGene 4.0, an extensible Java QTL-analysis platform. Bioinformatics. 2008;24:2788–9.

57. Howe KL, Achuthan P, Allen J, Allen J, Alvarez-Jarreta J, Amode MR, et al. Ensembl 2021. Nucleic Acids Research. 2021;49:D884–91.

58. Wang K, Li H, Xu Y, Shao Q, Yi J, Wang R, et al. MFEprimer-3.0: quality control for PCR primers. Nucleic Acids Research. 2019;47:W610–3.

59. Bolger AM, Lohse M, Usadel B. Trimmomatic: a flexible trimmer for Illumina sequence data. Bioinformatics. 2014;30:2114–20.

60. Eagle J, Ruff T, Hooker M, Sthapit S, Marston E, Marlowe K, et al. Genotyping by Multiplexed Sequencing (GMS) protocol in Barley. Euphytica. 2021;217:77.

61. Chen S, Zhou Y, Chen Y, Gu J. fastp: an ultra-fast all-in-one FASTQ preprocessor. Bioinformatics. 2018;34:i884–90.

62. Koren S, Walenz BP, Berlin K, Miller JR, Bergman NH, Phillippy AM. Canu: scalable and accurate long-read assembly via adaptive k-mer weighting and repeat separation. Genome Research. 2017;27:722–36.

63. Katoh K, Standley DM. MAFFT Multiple Sequence Alignment Software Version 7: Improvements in Performance and Usability. Molecular Biology and Evolution. 2013;30:772–80.

64. Xu W, Tucker JR, Bekele WA, You FM, Fu Y-B, Khanal R, et al. Genome Assembly of the Canadian Two-row Malting Barley Cultivar AAC Synergy. G3 Genes|Genomes|Genetics. 2021;11:jkab031.

65. Hu H, Wang P, Angessa TT, Zhang X-Q, Chalmers KJ, Zhou G, et al. Genomic signatures of barley breeding for environmental adaptation to the new continents. Plant Biotechnology Journal. 2023;21:1719–21.

66. Sakkour A, Mascher M, Himmelbach A, Haberer G, Lux T, Spannagl M, et al. Chromosome-scale assembly of barley Cv. ‘Haruna Nijo’ as a resource for barley genetics. DNA Research. 2022;29:dsac001.

67. Zeng X, Xu T, Ling Z, Wang Y, Li X, Xu S, et al. An improved high-quality genome assembly and annotation of Tibetan hulless barley. Scientific Data. 2020;7:139.

68. Dai F, Wang X, Zhang X-Q, Chen Z, Nevo E, Jin G, et al. Assembly and analysis of a qingke reference genome demonstrate its close genetic relation to modern cultivated barley. Plant Biotechnology Journal. 2018;16:760–70.

69. Sato K, Mascher M, Himmelbach A, Haberer G, Spannagl M, Stein N. Chromosome-scale assembly of wild barley accession “OUH602.” G3 Genes|Genomes|Genetics. 2021;11:jkab244.

70. Jiang C, Lei M, Guo Y, Gao G, Shi L, Jin Y, et al. A reference-guided TILLING by amplicon-sequencing platform supports forward and reverse genetics in barley. Plant Communications. 2022;3:100317.

71. RAxML version 8: a tool for phylogenetic analysis and post-analysis of large phylogenies | Bioinformatics | Oxford Academic. https://academic.oup.com/bioinformatics/article/30/9/1312/238053. Accessed 1 Mar 2024.

72. Darling ACE, Mau B, Blattner FR, Perna NT. Mauve: multiple alignment of conserved genomic sequence with rearrangements. Genome research. 2004;14:1394–403.

